# Comparative activity of Ceftriaxone, Ciprofloxacin and Gentamicin as a function of bacterial growth rate probed by *Escherichia coli* chromosome replication in the mouse peritonitis model

**DOI:** 10.1101/438663

**Authors:** Maria Schei Haugan, Anders Løbner-Olesen, Niels Frimodt-Møller

## Abstract

Commonly used antibiotics exert their effect predominantly on rapidly growing bacterial cells, yet growth dynamics taking place during infection in a complex host environment remain largely unknown. Hence, means to measure *in situ* bacterial growth rate is essential to predict the outcome of antibacterial treatment. We have recently validated chromosome replication as readout for *in situ* bacterial growth rate during *Escherichia coli* infection in the mouse peritonitis model. By the use of two complementary methods (qPCR and fluorescence microscopy) for differential genome origin and terminus copy number quantification, we demonstrated the ability to track bacterial growth rate, both on a population average and on a single-cell level; from one single biological specimen. Here, we asked whether the *in situ* growth rate could predict antibiotic treatment effect during infection in the same model. Parallel *in vitro* growth experiments were conducted as *proof-of-concept*. Our data demonstrate that the activity of commonly used antibiotics Ceftriaxone and Gentamicin correlated with pre-treatment bacterial growth rate; both drugs performing better during rapid growth than during slow growth. Conversely, Ciprofloxacin was less sensitive to bacterial growth rate, both in a homogenous *in vitro* bacterial population and in a more heterogeneous *in vivo* bacterial population. The method serves as a platform to test any antibiotic’s dependency upon active *in situ* bacterial growth. Improved insight into this relationship *in vivo* could ultimately prove helpful in evaluating future antibacterial strategies.

**Importance:** Most antibiotics in clinical use exert their effect predominantly on rapidly growing bacterial cells, yet there is a lack of insight into bacterial growth dynamics taking place during infection *in vivo.* We have applied inexpensive and easily accessible methods for extraction of *in situ* bacterial growth rate from bacterial chromosome replication during experimental murine infection. This approach not only allows for a better understanding of bacterial growth dynamics taking place during the course of infection, but also serves as a platform to test the activity of different antibiotics as a function of pre-treatment *in situ* growth rate. The method has the advantage that bacterial growth rate can be probed from a single biological sample, with the potential for extension into clinical use in pre-treatment infected biological specimens. A better understanding of commonly used antibiotics’ level of dependency upon bacterial growth, combined with measurements of *in situ* bacterial growth rate in infected clinical specimens, could prove helpful in evaluating future antibacterial treatment regimens.

## Introduction

Studies of antibacterial activity are largely based on laboratory models, where balanced bacterial populations propagate rapidly in well-defined batch cultures with a finite quantity of life-sustaining nutrients. These *in vitro* models fail to mirror the true growth dynamics of bacterial pathogens taking place during infection *in vivo*, where growth in a bacterial population appears to be both slower and less homogeneous (1–3). The dependency upon active bacterial growth for most antibiotics to exert their effect is acknowledged and has been examined by other investigators, both *in vitro* and *in vivo* (4–11). However, these studies were largely based on bacterial count kinetics as a direct measure of bacterial growth rate. This measure could be misleading during infection *in vivo*, where the net change in bacterial count is a function not only of growth, but also of elimination of bacterial cells by the host immune system; a factor not taken into account in the bacterial count kinetics method. Moreover, the method fails to report on any growth heterogeneity within the bacterial population. Hence, there is a need for refined means to measure bacterial growth that can extend into clinical use. To date, no gold standard method for measuring *in vivo* bacterial growth rate exists. In recent years, it has been demonstrated possible to extract direct measures of *in vivo* bacterial growth rate by analysing differential genome coverage in whole-genome sequencing data (12, 13). This method circumvents the limitation of a non-quantifiable bacterial elimination factor, as it reports directly on the bacteria’s physiological state. Nonetheless, it reports merely on the population mean growth rate.

Most bacterial chromosomes are circular with a single origin of replication (*oriC*), from where chromosome replication is initiated and carried out bidirectionally toward a single, opposite located terminus (*terC*) during bacterial growth (14, 15). In *Escherichia coli* it is acknowledged, from decades of *in vitro* studies, that bacterial growth rate is a function of growth conditions and is precisely coordinated with genome replication (16–18). When growth conditions are favourable, overlapping rounds of synchronously initiated bidirectional chromosome replication occur, allowing for the presence of more than two *oriC*s (number of *oriC* copies = 2^n^ (n = 2, 3, 4)) in rapidly growing cells (17, 19). In contrast, when growth conditions are disadvantageous, no or merely one round of chromosome replication occur, allowing for the presence of only one or maximum two *oriC*s (number of *oriC* copies = 2^n^ (n = 0, 1)) in non-or slowly growing cells (19). Hence, the copy number ratio of *oriC* to *terC* (i.e. *ori:ter*) reflects the bacterial population growth rate: during rapid growth larger fractions of cells undergo one or more round(s) of chromosome replication (i.e. population mean *ori:ter* ≥ 2) and during slow or no growth only few cells will undergo chromosome replication (i.e. population mean *ori:ter* ~ 1) (20). By the use of two complementary methods for measuring *ori:ter*: quantitative PCR (qPCR) and fluorescence microscopy, we have been able to probe *in situ* growth rates of fluorescently labelled *E. coli* ATCC^®^ 25922™ both on a population average (by qPCR) and on a single-cell (by fluorescence microscopy) level during wide-spread infection in the mouse peritonitis model (3). We demonstrated correlation between *ori:ter* and bacterial cell size at all growth rates and the ability of *ori:ter* to predict the development in net bacterial population size. Moreover, in this recent observation of growth dynamics during host infection we found that growth rates were largely heterogeneous within the bacterial populations propagating both in the peritoneum and in the blood, respectively, throughout the length of infection (3). This finding is in consistency with previous reports of *Staphylococcus aureus* growth rate heterogeneity in cystic fibrosis sputum, as measured by isotope tracing (1). These observations underscore the need for refined and easily accessible methods to measure the *in situ* bacterial growth rate taking place during various types of infection and its causal relationship with the outcome of antibacterial treatment.

Here, we extended the approach of using chromosome replication as readout for *in vivo* bacterial growth rate in the mouse peritonitis model to explore its potential in predicting antibacterial treatment effect. For comparison, we chose a representative drug from each of three classes of commonly used bactericidal antibiotics with different cellular targets: Ceftriaxone (CRO; a β-lactam), Ciprofloxacin (CIP; a Fluoroquinolone) and Gentamicin (GEN; an Aminoglycoside) (21). We hypothesized that the drugs tested would perform better when given during rapid, than during slow bacterial growth.

## Results

We defined the minimal and maximal growth rates of *E. coli* during infection in the mouse peritonitis model and compared the activity of standardised antibacterial dosing regimens given during either rapid or slow bacterial growth, as defined by qPCR-derived *ori:ter* from infected body fluids. As qualitative control, fluorescence microscopy was applied on the same materials to demonstrate any treatment-induced physiological change in cell morphology or chromosome replication on a single-cell level. As *proof-of-principle*, corresponding *in vitro* treatment experiments using the same model infective organism, the EUCAST and CLSI reference strain, *E.coli* ATCC^®^ 25922™, were carried out.

### Antibacterial activity as a function of growth rate *in vitro*

*E. coli* growth experiments in batch cultures were performed to validate the methods applied in the experimental *in vivo* infection model. For these experiments we used Lysogeny Broth (LB), a rich media that supports rapid bacterial growth, allowing cells to reach stationary phase due to exhaustion of utilisable carbon sources before any significant physiological alterations in growth media occur (22). Here, stationary phase bacterial cells were diluted in fresh media and allowed to grow with repeated sample collections at 2, 4, 8 and 10 hours of incubation. All antibiotics were given as a single-dose during rapid or slow bacterial growth, respectively. Antibiotic concentrations were standardised and defined from previous studies, and corresponded to serum concentrations observed in humans on standard dosing regimens: Ceftriaxone 30 mg/l; Ciprofloxacin 1 mg/l; Gentamicin 10 mg/l (23–28). Sampling was performed after two hours of antibiotic exposure. There was temporal difference in pre-treatment bacterial load at the time points chosen for rapid and slow bacterial growth treatment induction, respectively. Hence, antibiotic activity was measured as the difference in pre- and post-treatment bacterial count, relative to the pre-treatment bacterial count (i.e. relative Δlog_10_ CFU/ml), for comparison. We tested the wild-type bacterial strain (ATCC 25922) in parallel to the genetically modified derivative of the strain, with chromosomally incorporated fluorescent *oriC* and *terC* labels (ALO 4783), to ensure the absence of growth retardation due to transgene insertion. As no growth differences were observed (Fig. 1a), *in vitro* data from both versions of the strain were pooled for analysis. In regard to the Gentamicin treatment regimen, however, only wild-type ATCC 25922 was applied, as the Gentamicin MIC was increased by the presence of the non-removable Kanamycin cassette encoding Kanamycin phosphotransferase, used as clonal selection marker, in ALO 4783 (3, 29) (Table 1).

**FIG 1.**
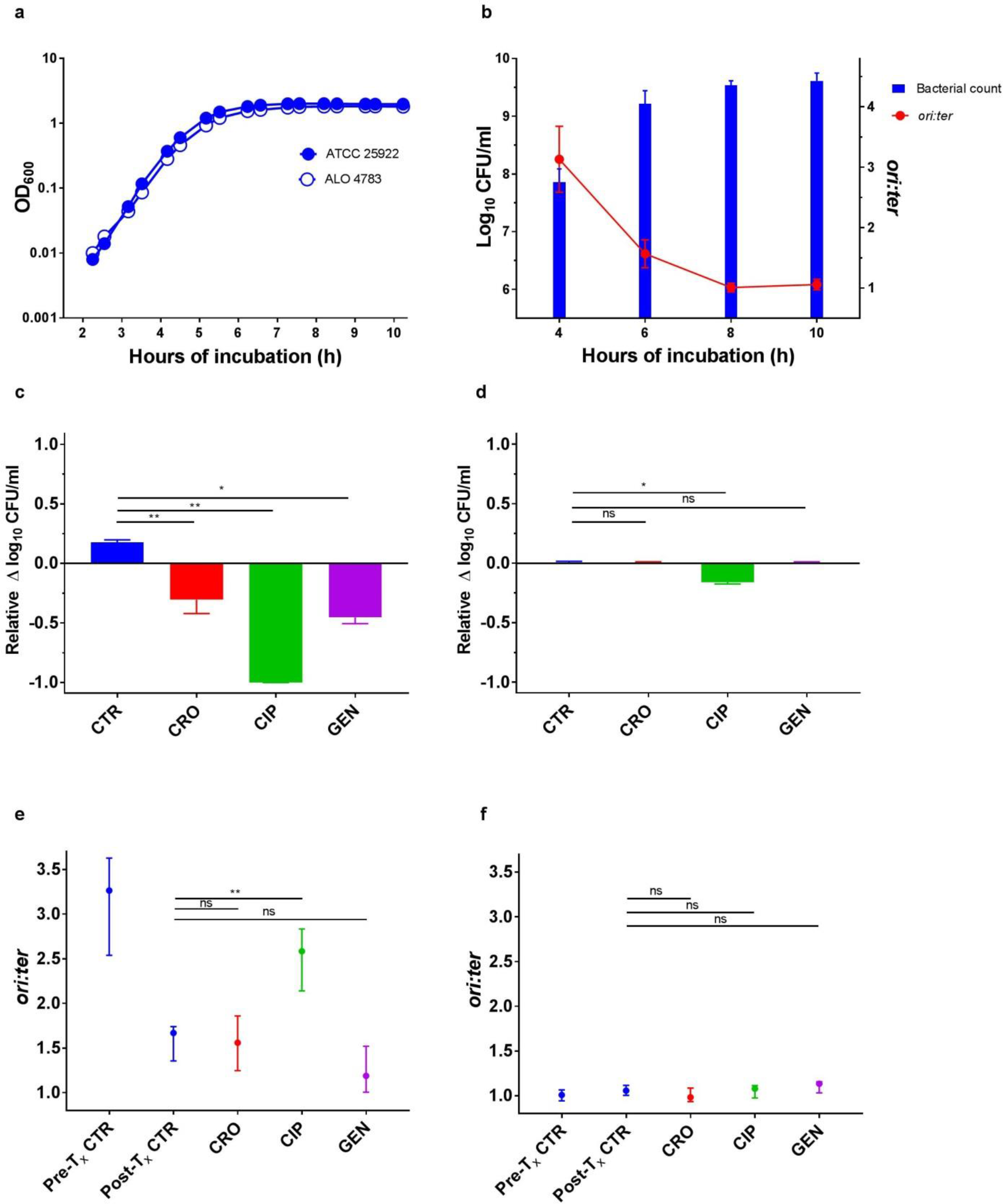
Antibiotic activity as a function of bacterial growth rate *in vitro* **(a)** Parallel growth of ALO 4783 relative to ATCC 25922 wild-type in LB revealed no growth retardation due to transgene insertions. Cell density measured as OD_600_. **(b)** Bacterial counts (CFU/ml) and growth rates (*ori:ter*) in untreated control batch cultures (ATCC 25922 and ALO 4783). n = 6. Data presented as mean (SD) **(c)** Bacterial count reductions after two hours of antibiotic exposure in Ceftriaxone (CRO), Ciprofloxacin (CIP) and Gentamicin (GEN) treatment batch cultures when therapy was induced during rapid bacterial growth (i.e. at 4 hours of incubation). Controls (CTR) received no antibiotic therapy. **(d)** Bacterial count reductions after two hours of antibiotic exposure in treatment batch cultures when therapy was induced during slow bacterial growth (i.e. at 8 hours of incubation). CTRs received no antibiotic therapy. For comparison of activity between treatment induction during rapid and slow growth, respectively, data in **(c)** and **(d)** are presented as relative bacterial count reductions. **(e)** Bacterial growth rates (*ori:ter*) in pre- and post-treatment controls and in treatment batch cultures after two hours of antibiotic exposure when therapy was induced during rapid bacterial growth. **(f)** Bacterial growth rates (*ori:ter*) in pre- and post-treatment controls and in treatment batch cultures after two hours of antibiotic exposure when therapy was induced during slow bacterial growth. Data in **(c)** – **(f)** are presented as median and interquartile range (IQR). CTR, n = 6; CRO, n = 6; CIP, n = 6; GEN, n = 3. *P* by Mann-Whitney U test (*, *P* < 0.05; **, *P* < 0.01; ns, *P* > 0.05).

**TABLE 1.**
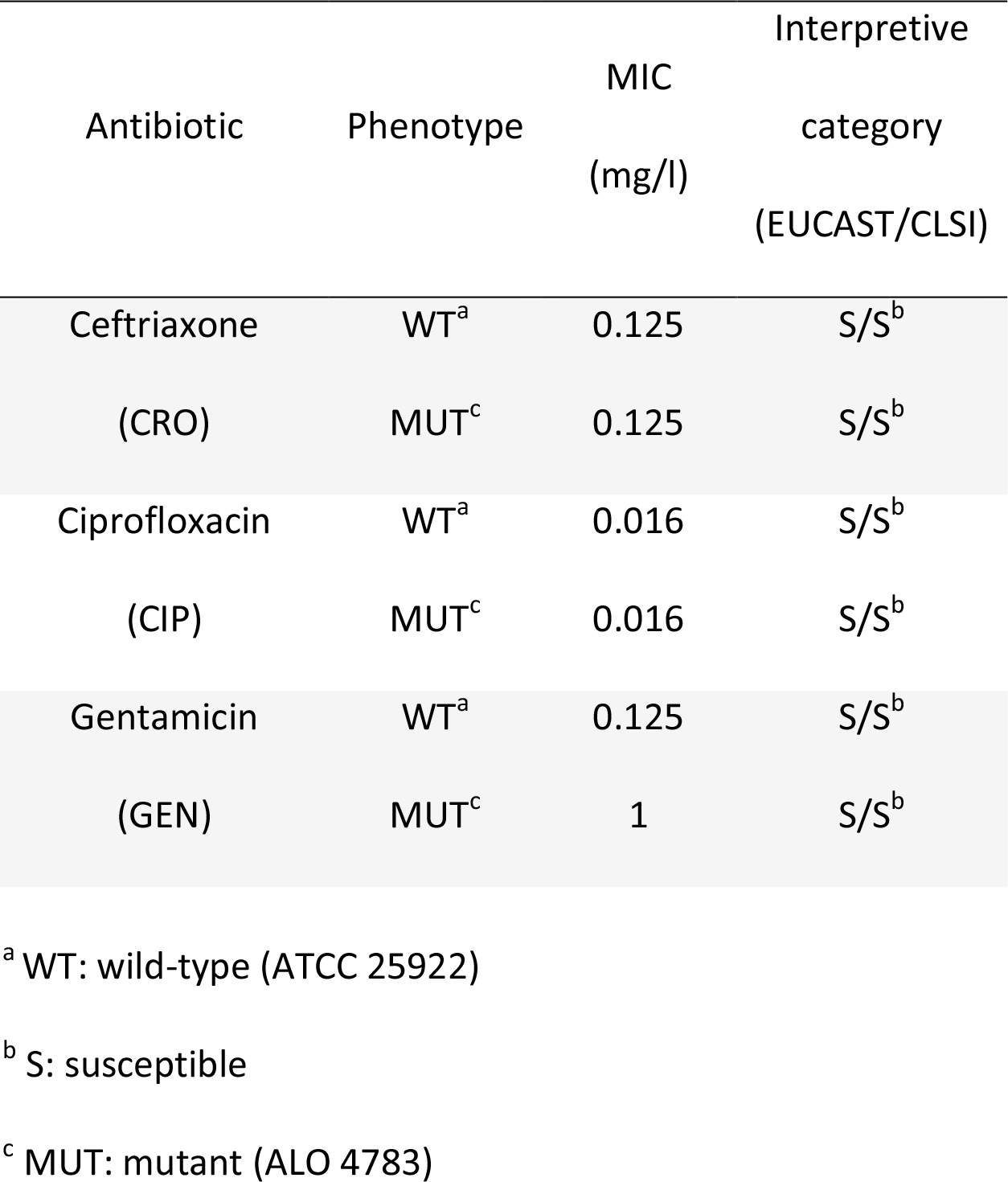
Overview of MIC and susceptibility interpretation of the two isogenic strains applied in *in vitro* and *in vivo* experiments

### Antibiotics administered during rapid bacterial growth *in vitro*

The bacterial populations propagating in batch cultures reached maximal growth rates at 4 hours of incubation (expressed as mean (SD) *ori:ter* = 3.13 (0.55)) (Fig. 1b). At this stage, it has been demonstrated that *E. coli* population growth in LB is close to balanced and dominated by large cells growing with overlapping rounds of chromosome replication (3). This is exemplified in Figure 2, image A, illustrating a large bacterial cell with multiple *oriC*s. As a consequence of rapid bacterial growth there was subsequent increase in population size (Fig. 1a – b). Between 4 and 6 hours of incubation growth was starting to slow down, illustrated by a reduction of *ori:ter* at 6 hours of incubation and following minimal increase in net population size (Fig. 1a – b).

**FIG 2.**
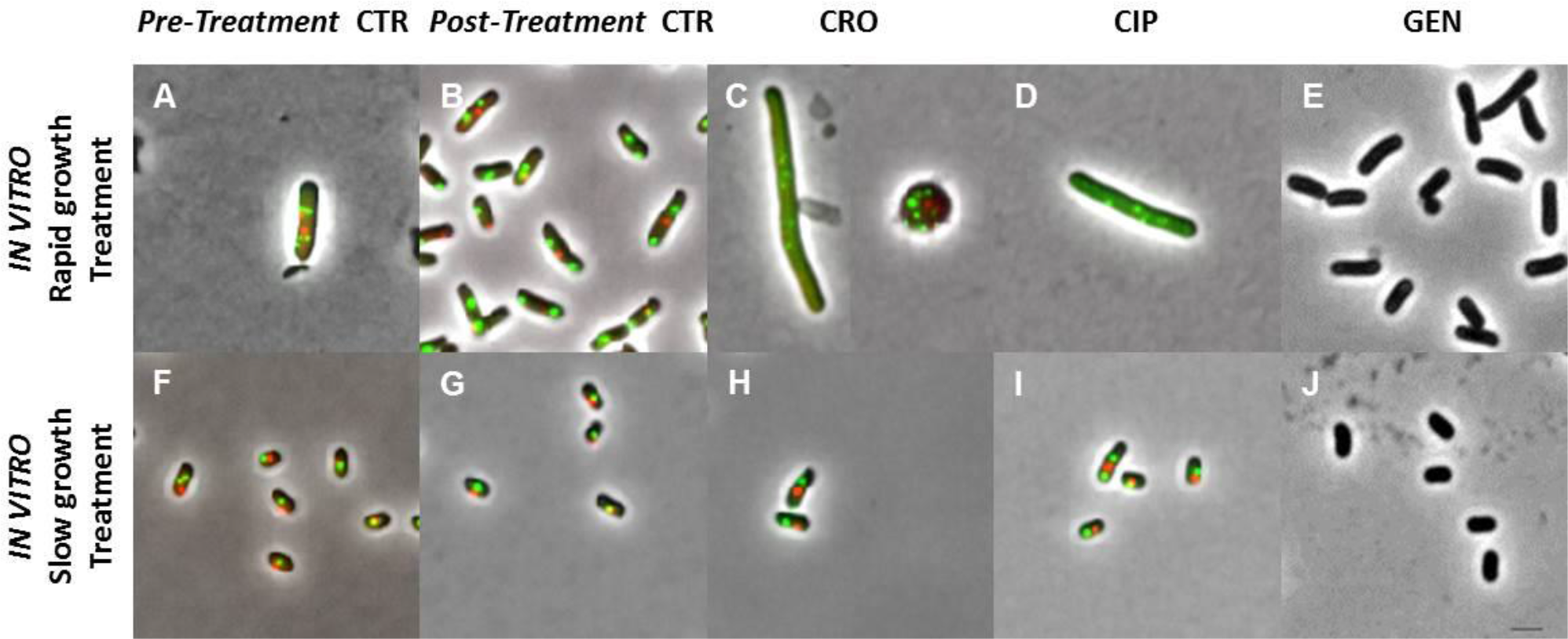
Representative examples of pooled bacterial cells isolated by fluorescence microscopy after antibiotic induction during rapid (top row) or slow bacterial growth (bottom row) *in vitro.* Images are shown in phase contrast; intracellular *oriC* foci in green (GFP) and *terC* foci in red (mCherry) (ALO 4783). For GEN treatment experiments ATCC 25922 without fluorescence foci was applied. A total of n = 500 pooled cells were analysed per time point from all cultures with the exception of rapid growth treatment cultures: CRO, n = 28; CIP, n = 112; GEN, n = 330. Due to limitation in fluorescence microscopy resolution of co-localising *oriC*s, some bacterial cells with overlapping chromosome replication might appear with too few foci (33). Population mean (SD) medial axis cell lengths (μm) were as follows: A: 4.1 (0.98); B: 3.51 (0.86); C: not determined (due to overrepresentation of spherical cells); D: 7.39 (2.52); E: 3.25 (0.84); F: 2.58 (0.66); G: 2.25 (0.52); H: 2.67 (0.67); I: 2.53 (0.69); J: 2.18 (0.45). CTR: control; CRO: Ceftriaxone; CIP: Ciprofloxacin; GEN: Gentamicin. Scale bar is 2 μm.

When introduced into a batch culture of rapidly growing cells, all antibiotics resulted in significant bacterial count reduction, compared to controls (CRO and CIP, *P* < 0.001; GEN, *P* < 0.01) (Fig. 1c). Correspondingly, single-cell microscopic analysis of pooled live bacterial cells from treatment batch cultures revealed that the majority of the cells we were able to isolate after exposure to either Ceftriaxone or Ciprofloxacin were affected by the treatment, when introduced during rapid bacterial growth (Fig. 2, image C and D, respectively). These bacterial populations were represented predominantly by spherical and filamentous cells (Ceftriaxone), or elongated cells (Ciprofloxacin); all with multiple fluorescent foci, representing direct or indirect chromosome replication disturbance. Unfortunately, we were unable to accurately quantify the population distribution of *oriC and terC*, respectively, due to multiple, overlapping fluorescent foci in these treatment groups. The morphological changes in the Ceftriaxone exposed bacterial populations suggest that Ceftriaxone exerted its bactericidal effect through cell-wall inhibition predominantly by binding to Penicillin-binding proteins (PBPs) 2 and 3, in accordance with previous studies (30). We underscore that the microscopy data from bacterial cultures treated with antibiotics during rapid growth (Fig. 2, image C - E) are subject to uncertainty given that only few live bacterial cells (n < 500) were isolated due to the efficient bacterial killing (Fig. 1c). Nevertheless, *ori:ter* extracted from qPCR complemented the above described microscopy findings regarding chromosome replication: high *ori:ter* ratios were observed where photomicrographs were dominated by large cells with multiple *oriCs*, and low *ori:ter* ratios where photomicrographs were dominated by small cells with few *oriCs*, in both treatment and control groups (Fig. 1e and Fig. 2, images A – E). Only Ciprofloxacin administered during rapid growth entailed significantly higher *ori:ter*, compared to post-treatment controls (*P* < 0.01) (Fig. 1e). As to Ceftriaxone and Gentamicin treatment administered during rapid bacterial growth, both induced a decrease in *ori:ter* toward ~ 1, similar to that of the control group (Fig. 1e). For Gentamicin treated bacterial cells, microscopic visualisation of *oriC* and *terC* was not possible as only the wild-type ATCC 25922 was applied. The size and morphology of the cells did, however, not appear to differ from that of the respective control population (Fig. 2, images E and B, respectively).

### Antibiotics administered during slow bacterial growth *in vitro*

At 8 hours of incubation minimal bacterial growth rates (expressed as mean (SD) *ori:ter* = 1.01 (0.07)) were reached due to nutrient starvation (Fig. 1b). At this stage, it has been demonstrated that the bacterial population is dominated by small bacterial cells without ongoing chromosome replication, i.e. complete or near complete cessation of growth (3). This is exemplified in Figure 2, image F, illustrating small bacterial cells with predominantly one *oriC*/cell. Consequently, there was no significant subsequent net population size increase, and all parameters remained largely unchanged by 10 hours of incubation (Fig. 1a – b and Fig. 2, image G). When identical antibiotic regimens as those applied during rapid bacterial growth were introduced into this population of slowly / non-growing bacterial cells, only Ciprofloxacin caused significant bacterial count reduction (*P* < 0.01); albeit considerably less than when administered during rapid growth, where a near total clearance of cells was observed (Fig. 1c - d). Ceftriaxone and Gentamicin treatment effect was absent (Fig. 1d). Correspondingly, photomicrographs were dominated by cells largely unaffected by all three antibiotics, when compared to post-treatment controls (Fig. 2, images G – J), and no significant change in *ori:ter* was observed in any treatment group, compared to post-treatment controls (Fig. 1f). The Ciprofloxacin-induced increase in *ori:ter* observed during rapid bacterial growth treatment (Fig. 1e and Fig. 2, image D) was absent when Ciprofloxacin was added to a population of cells largely without ongoing chromosome replication (mean population *ori:ter* ~ 1) (Fig. 1f and Fig. 2, image I).

In summary, during controlled bacterial growth in a closed, rich media batch culture extreme situations of both rapid growth and complete or near complete cessation of growth could be provoked. When identical antibiotic treatment regimens were introduced to cultures of bacterial populations growing at either maximal or minimal growth rate, the dependency of active bacterial growth for all drugs to exert their effect became evident. The significant bacterial load reductions observed when Ceftriaxone or Gentamicin was added to rapidly growing bacterial populations, were lost when identical treatment regimens were induced during slow bacterial growth. Ciprofloxacin, however, was less sensitive to active bacterial growth, as a significant reduction in bacterial load was observed upon administration during both rapid and slow growth.

### Antibacterial activity as a function of growth rate *in vivo*

In the experimental mouse peritonitis model a total of 54 mice pooled from 4 independent experiments were challenged intraperitoneally with stationary phase *E. coli*. All animals developed wide-spread infection within 2 hours post challenge. Maximal and minimal bacterial growth rates were successfully probed from infected body fluids (peritoneal lavage fluid (PLF) and blood) (3).

During propagation in this infection model, bacterial growth was overall slower than *in vitro*, yet never came to a complete cessation (i.e. *ori:ter* remained > 1 at all times) during the length of the experiment (Fig. 3a). All antibiotics were administered as a single-dose subcutaneously (s.c.) during rapid or slow bacterial growth, respectively, in the following concentrations: Ceftriaxone 5 mg, Ciprofloxacin 0.4 mg and Gentamicin 1 mg per mouse, respectively. Antibiotic concentrations were defined from previous studies (23–28) and all were > minimum 10×MIC (Table 1). Infected biological specimens (PLF, blood, spleen and kidneys) were harvested after 2 hours of antibiotic exposure. For Ceftriaxone and Ciprofloxacin treatment experiments, ALO 4783 was used as infective agent. For Gentamicin treatment experiments ATCC 25922 wild-type was applied, for reasons previously explained.

**FIG 3.**
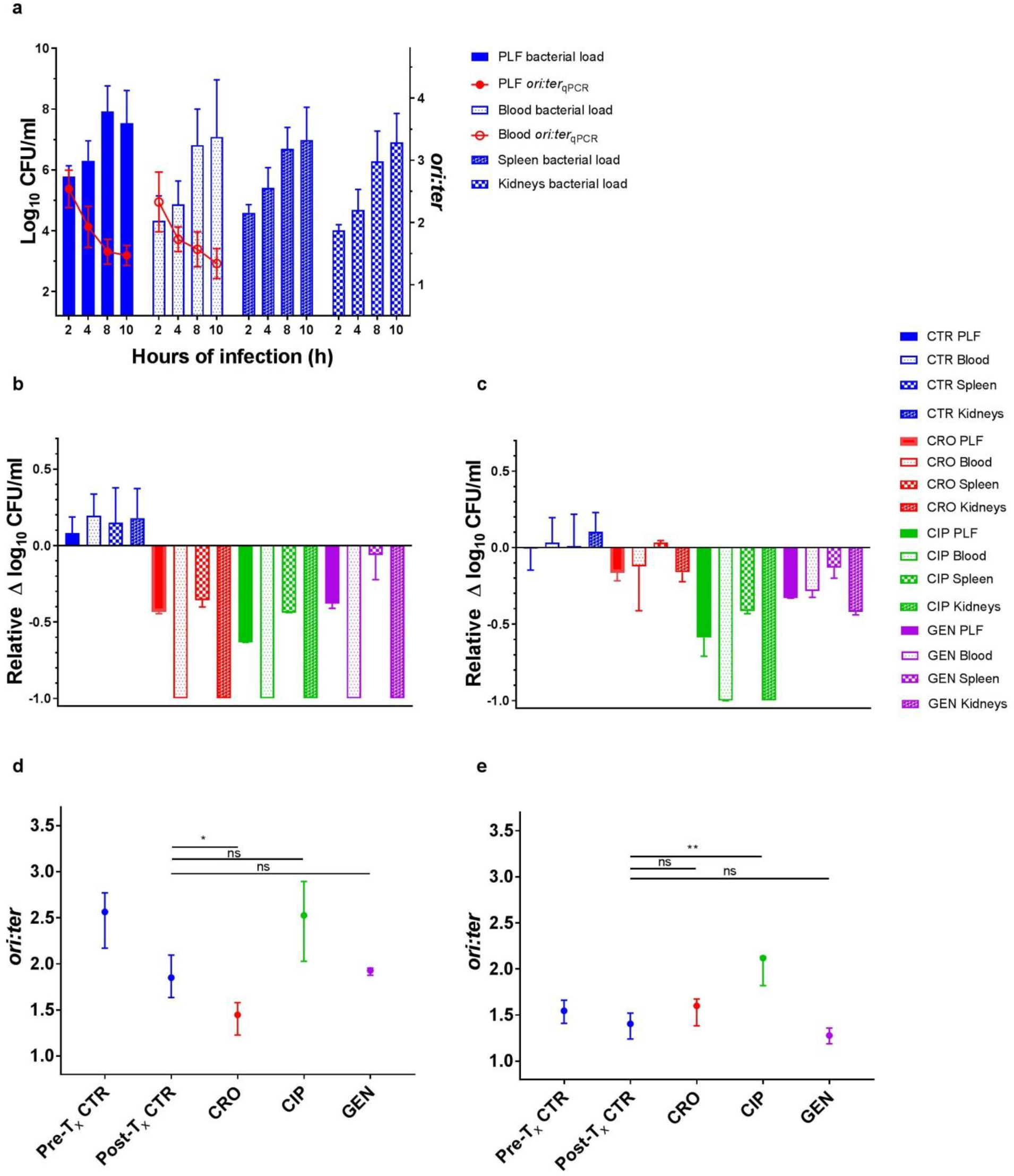
Antibiotic activity as a function of bacterial growth rate during infection *in vivo* in the mouse peritonitis model **(a)** Bacterial counts (CFU/ml; n = 9 per time point) and growth rates (*ori:ter;* 2, 8 and 10 hours of infection, n = 9; 4 hours of infection, n = 6) in untreated control groups (ATCC 25922 and ALO 4783) in the peritoneal lavage fluid (PLF), blood, spleen & kidneys (bacterial counts only in the tissues). **(b)** Bacterial count reductions in PLF, blood, spleen and kidneys after two hours of antibiotic exposure in Ceftriaxone (CRO), Ciprofloxacin (CIP) and Gentamicin (GEN) treatment groups when therapy was induced during rapid bacterial growth (i.e. at 2 hours of infection). Controls (CTR) received no antibiotic therapy. **(c)** Bacterial count reductions in PLF, blood, spleen and kidneys after two hours of antibiotic exposure in treatment groups when therapy was induced during slow bacterial growth (i.e. at 8 hours of infection). CTRs received no antibiotic therapy. For comparison of activity between treatment induction during rapid and slow growth, respectively, data in **(b)** and **(c)** are presented as relative bacterial count reductions. CTR, n = 9; CRO, CIP and GEN, n = 3 **(d)** Bacterial growth rates (*ori:ter*) in pre- and post-treatment (T_x_) controls and in treatment groups after two hours of antibiotic exposure when treatment was induced during rapid bacterial growth. As there was no significant difference in *ori:ter* between control bacterial populations from PLF and blood, these were pooled for analysis. *ori:ter* in treatment groups were only available from PLF, due to total bacterial elimination from the blood. Pre-treatment CTR n = 18; Post-treatment CTR, n = 12; CRO, n = 3; CIP, n = 3; GEN, n = 3. **(e)** Bacterial growth rates (*ori:ter*) in pre- and post-treatment (T_x_) controls and in treatment groups after two hours of antibiotic exposure when treatment was induced during slow bacterial growth. PLF and blood *ori:ter* were pooled for analysis. Pre-treatment CTR n = 18; Post-treatment CTR, n = 18; CRO, n = 6; CIP, n = 3; GEN, n = 6. Data in **(b) – (e)** are presented as median and interquartile range (IQR). *P* by Mann-Whitney *U* test (*, *P* < 0.05; **, *P* < 0.01; ns, *P* > 0.05).

### Antibiotics administered during rapid bacterial growth *in vivo*

The bacterial populations reached maximal *in situ* growth rates (expressed as mean (SD) *ori:ter* PLF = 2.54 (0.30); *ori:ter* blood = 2.37 (0.46)) at 2 hours of infection (Fig. 3a). At this stage of infection, it has been demonstrated that bacterial population growth is heterogeneous (i.e. the population is made up of bacterial cells of various sizes and DNA content) (3). Figure 4, image A, illustrates a representative large cell with multiple *oriC*s, isolated from the PLF. As a consequence of average rapid growth rates, there was subsequent increase in net bacterial population size in all biological specimens (Fig. 3a). After 2 hours of infection there was a gradual decrease in *ori:ter*, resulting in overall net population stagnation between 8 and 10 hours of infection (Fig. 3a).

**FIG 4.**
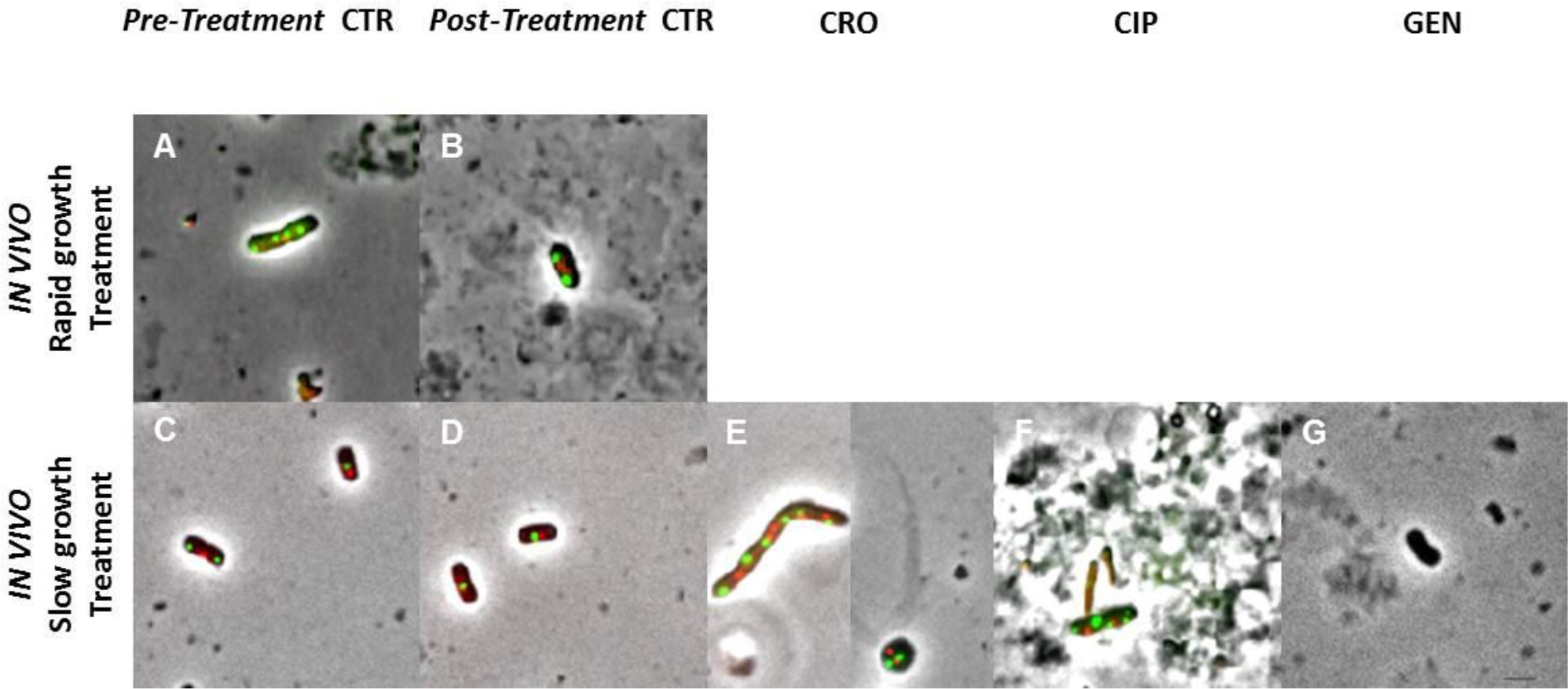
Representative examples of bacterial cells isolated by fluorescence microscopy after antibiotic induction during rapid (top row) or slow bacterial growth (bottom row) *in vivo* (blood and peritoneal lavage fluid (PLF) bacterial cells pooled). Images are shown in phase contrast; intracellular *oriC* foci in green (GFP) and *terC* foci in red (mCherry) (ALO 4783). For GEN treatment experiments ATCC 25922 without fluorescence foci was applied. Examples shown are bacteria isolated from the PLF. Due to total or near total elimination of bacterial cells during rapid growth treatment, no bacterial cells were available from these treatment groups. A total of n = 500 cells pooled were analysed per time point from pre- and post-treatment controls during slow growth, respectively. For pre- and post-treatment controls during rapid growth, fewer cells were isolated due to low bacterial counts during early hours of infection: n = 142 and n = 66, respectively. For slow growth treatment induction: CRO, n = 170; CIP, n = 35; GEN, n = 228. Due to limitation in fluorescence microscopy resolution of co-localising *oriC*s, some bacterial cells with overlapping chromosome replication might appear with too few foci (33). Population mean (SD) medial axis cell lengths (μm) were as follows: A: 3.96 (1.15); B: 3.73 (1.13); C: 3.19 (0.73); D: 2.77 (0.86); E: not determined (due to overrepresentation of spherical cells); F: 3.25 (0.68); G: 2.56 (0.76). CTR: control; CRO: Ceftriaxone; CIP: Ciprofloxacin; GEN: Gentamicin. Scale bar is 2 μm.

When administered during rapid bacterial growth, all antibiotics (CRO, CIP and GEN) caused significant bacterial count reduction at all anatomical sites examined (PLF, blood, spleen and kidneys, *P* < 0.01), including a total elimination in blood and kidneys, compared to respective controls (Fig. 3b). The anatomical site-specific differences in antibiotic activity observed (e.g. between the blood and PLF) (Fig. 3b) cannot be explained by differences in site-specific *in situ* pre-treatment *ori:ter*, as these were not significantly different (*P* > 0.5) (Fig. 3a). Rather, these differences are conceivably attributable to other pharmacodynamic and / or host immune parameters.

We were able to isolate only few live bacterial cells for fluorescence microscopy from infected body fluids in pre- and post-treatment control groups during rapid bacterial growth (Fig. 4, images A - B) due to low bacterial counts during early hours of infection (Fig. 3a). Given the substantial or total clearance of bacteria from the PLF and blood following antibiotic exposure during rapid bacterial growth, we were unable to isolate live bacterial cells from these treatment groups (Fig. 4). However, the overall *ori:ter* developments in these treatment groups were similar to those observed during rapid growth *in vitro*: Ciprofloxacin treatment induced significantly higher *ori:ter* ratios compared to post-treatment controls (*P* < 0.05), and *ori:ter* developments in Ceftriaxone and Gentamicin treatment groups were similar to those of the post-treatment controls (i.e. reduction toward ~ 1); with Ceftriaxone being most efficient (*P* < 0.05) (Fig. 3d).

### Antibiotics administered during slow bacterial growth *in vivo*

Minimal bacterial growth rates (expressed as mean (SD) *ori:ter* PLF = 1.53 (0.2) and *ori:ter* blood = 1.57 (0.29)) were observed starting from 8 hours of infection (Fig. 3a). Marginally lower *ori:ter* levels were observed in bacteria isolated from both PLF and blood at 10 hours of infection (mean (SD) PLF = 1.48 (0.17) and *ori:ter* blood = 1.34 (0.24)) (Fig. 3a). However, these differences were not statistically significant (*P* > 0.5), and, at this time the criteria for euthanasia were met for control animals. Hence, 8 hours of infection was chosen as the time point for slow bacterial growth treatment induction.

When administered during slow bacterial growth, only Ciprofloxacin treatment caused significant bacterial count reductions at all anatomical sites (PLF, blood, spleen and kidneys, *P* < 0.01 (Fig. 3c). The activity of Gentamicin was overall reduced, and Ceftriaxone activity was substantially reduced at all anatomical sites when antibiotics were administered during slow bacterial growth, compared to during rapid bacterial growth (Fig. 3b - c).

Photomicrographs of live bacterial cells exposed to antibiotics during slow bacterial growth indicated that the drugs exerted their effect similar to that *in vitro*: Ceftriaxone inhibiting cell wall synthesis predominantly via PBP 2 (demonstrated by the presence of spherical cells) and PBP 3 (demonstrated by the presence of filamentous cells) (30) (Fig. 4, image E), and Ciprofloxacin interrupting natural bacterial growth by inducing cell enlargement with multiple fluorescent foci due to interference with ongoing chromosome replication (Fig. 4, image F) (31). Due to multiple, overlapping fluorescent foci in these treatment groups, we were unable to accurately quantify the population distribution of *oriC and terC*, respectively. For Gentamicin treated bacterial cells, microscopic visualisation of *oriC* and *terC* was not possible as the wild-type ATCC 25922 was applied. The size and morphology of the bacterial cells did, however, not appear to differ from that of the respective control population, as observed after antibiotic administration during slow bacterial growth (Fig. 4, Images D and G). We emphasize the uncertainty in these microscopy data (Fig. 4, Images H – J), as only few live bacteria (n < 500) were isolated due to antibiotic induced bacterial killing (Fig. 3c).

The bacterial populations (both in PLF and blood) exposed to antibiotics during late stage of infection (i.e. at 8 hours of infection, Fig. 3a) differed from that observed after prolonged propagation *in vitro* (i.e. at 8 hours of incubation; Fig 1b) in that there was no complete cessation of growth in the former. Here, fractions of the population were still undergoing chromosome replication (as expressed by the mean *ori:ter* > 1). Hence, the effect of Ciprofloxacin on *ori:ter* was apparent also upon treatment during slow bacterial growth (Fig. 3e).

In summary, *in vivo* bacterial growth rates at the time points representing rapid and slow bacterial growth, respectively, did not differ to the same extent as those during propagation in a rich media *in vitro.* Consequently, the difference in antibacterial activity as a function of bacterial growth rate became less explicit. Nevertheless, the overall trends were similar to those observed *in vitro:* only Ciprofloxacin treatment entailed significant bacterial load reduction in all examined body fluids and tissues, both during rapid and slow bacterial growth. Contrary to the *in vitro* results, however, Ceftriaxone and Gentamicin both caused a certain bacterial load reduction when administered during slow bacterial growth, albeit overall less than during rapid bacterial growth. This difference is likely due to the fact that bacterial growth at 8 hours of infection was not at a (near) complete arrest, as the *ori:ter* remained > 1.

## Discussion

In this study, we determined the activity of three commonly used bactericidal antibiotics with different antibacterial targets as a function of *in situ* bacterial growth rate, expressed as differential genome origin and terminus copy number quantification (*ori:ter)* by qPCR. We demonstrated that overall activities of both Ceftriaxone and Gentamicin were substantially lower when administered during slow bacterial growth, than during rapid bacterial growth *in vivo.* Contrarily, Ciprofloxacin was less sensitive to bacterial growth rate, as the overall activity remained largely unchanged when going from rapid to slow bacterial growth rate treatment induction *in vivo.* The findings of Ciprofloxacin being less sensitive to bacterial growth rate than β-lactams and Aminoglycosides, respectively, has been demonstrated by others, however with the limitation of bacterial growth rate being estimated from net bacterial population kinetics (9, 10). In the parallel *in vitro* experiments, the difference between rapid and slow bacterial growth rate was more explicit, including complete or near complete cessation of growth as the lower extreme growth rate. When administered during near cessation of bacterial growth, only Ciprofloxacin exerted significant bacterial load reduction, while Ceftriaxone and Gentamicin lost their effect, in agreement with previous observations where bacterial growth rates were extracted from population kinetics (4, 8). The increase in *ori:ter* observed after administration of Ciprofloxacin to populations of rapidly growing cells, both *in vivo* and *in vitro*, confirms the drug’s mode of action. Ciprofloxacin exerts its effect predominantly through DNA gyrase inhibition, which results in the formation of double-strand DNA breaks during chromosome replication, prohibiting the replication forks from reaching the terminus (i.e. the copy number of *oriC* relative to *terC* will be high) (31). As anticipated, this effect was lost when Ciprofloxacin was introduced into a population without ongoing chromosome replication during slow growth treatment *in vitro*; as opposed to the slowly growing bacterial populations *in vivo*, where fractions of cells were still undergoing chromosome replication. As to Ceftriaxone and Gentamicin, both drugs induced a decrease in *ori:ter* toward ~ 1 in rapidly growing bacterial populations, both *in vitro* and *in vivo.* Contrary to the respective post-treatment control bacterial populations, however, where similar reductions in *ori:ter* were observed, this decrease cannot be explained by natural reduction of bacterial growth rate due to population entrance into stationary phase (i.e. starvation of life-sustaining nutrients due to high population density), as bacterial counts were reduced during antibiotic treatment.

Rather, the Ceftriaxone or Gentamicin induced *ori:ter* reduction toward ~ 1 is likely the result of preferential elimination of fractions of rapidly growing bacterial cells (i.e. those with *ori:ter* > 1). Hence, besides allowing for measurement of pre-treatment *in situ* bacterial growth rate, *ori:ter* may to some extent, demonstrate antimicrobial mode of action by analysis of post-treatment *ori:ter*. Fluorescence microscopy can complement these findings by direct single-cell visualisation, as demonstrated, yet is limited by the absence of live bacterial cells after efficient bacterial elimination.

The growth rate scenarios observed during bacterial propagation *in vitro* were not representative of the bacterial growth dynamics taking place during infection in a complex host environment. In the latter, a complete or near complete cessation of bacterial growth was not observed as long as the host was alive and bacterial life-sustaining nutrients presumably not exhausted. Hence, for more meaningful prediction of antibiotic activity *in vivo,* it is important to be able to test this in relation to the *in situ* bacterial growths rate taking place during host infection, rather than extrapolating from *in vitro* studies. *ori:ter* provides predictive value in informing on the likelihood of antibiotic activity, both during *E. coli* propagation *in vitro* and *in vivo* during host infection. There are, however, limitations to be considered in this study. Maximal bacterial growth rates were observed after few hours of propagation, while minimal growth rates were only observed after prolonged propagation, both *in vitro* and *in vivo*. Consequently, the net bacterial population sizes were larger during slow, than during rapid bacterial growth. For meaningful comparison of effect between identical antibacterial treatment regimens applied, we calculated the relative killing effect in both scenarios. We cannot exclude the possibility of an inoculum effect as contributing factor to the lower antibiotic activity observed during slow bacterial growth. This is, however, a phenomenon mainly observed for β-lactams and rather unlikely to have occurred at the high antibiotic concentrations that were applied, both *in vitro* and *in vivo* (32). Moreover, we were unable to adequately purify bacterial DNA from spleen and kidney tissues. Hence, the difference in spleen and kidney antibiotic treatment effect between the two scenarios *in vivo* is only assumed to owe to different *in situ* bacterial growth rate in these tissues. Yet, as the temporal development in net bacterial population size from these tissues followed those of the PLF and blood, we find it likely that bacterial growth rates in the tissues would also be higher during early hours of infection, than after prolonged propagation.

We conclude that chromosome replication as a means to measure bacterial growth rate can predict antibacterial treatment outcome. To some extent it can also elucidate the antibiotic mode of action, as exemplified by the increase in *ori:ter* caused by Ciprofloxacin-induced double-strand DNA breaks, and the decrease in *ori:ter* caused by preferential elimination of rapidly growing bacterial cells by both Ceftriaxone and Gentamicin. While our findings are in agreement with previously demonstrated causal relationship between *in vivo* bacterial growth rate and antibiotic activity, previous studies were limited by the methodology; e.g. the bacterial populations kinetics method fails to take into account the host elimination factor, and tracking of bacterial growth by isotope trace incorporation is largely inconvenient when it comes to pursuing the method into clinical practice. Tracking bacterial growth rate by differential genome origin and terminus quantification by qPCR has the advantage of being accessible, inexpensive and reports directly on the bacterial physiology, circumventing the limitation of the bacterial count kinetics method. Also, growth rates can be probed from a single biological sample, which is convenient in a clinical setting where repeated sample measurement often is difficult. The method could serve as a platform for testing any antimicrobial’s activity as a function of pre-treatment bacterial growth rate in experimental infection models - and could be pursued in a clinical setting to examine bacterial growth rates in infected biological materials. This could in turn prove helpful in evaluating future antibacterial strategies.

## Materials and Methods

### Bacterial strains

*Escherichia coli* ATCC^®^ 25922™, a clinical isolate from the American Type Culture Collection (Manassas (VA), USA) and CLSI and EUCAST control strain for antibiotic susceptibility testing, was used throughout the study. This strain was applied both as wild-type and as a genetically modified version expressing fluorescent fusion-proteins at chromosomal sites corresponding to *oriC* and *terC,* respectively (ALO 4783) (3).

### Antimicrobial agents and susceptibility testing

Antimicrobial agents used in this study were procured as the commercial products registered for parenteral use in Denmark. Ciprofloxacin (CIP) as Ciprofloxaxin 2 mg/ml (Fresenius Kabi, Germany), Ceftriaxon (CRO) as Ceftriaxon “Stragen” 1 g (Stragen Nordic, Denmark) and Gentamicin (GEN) as Hexamycin, 40 mg/ml (Sandoz, Denmark). Ceftriaxone was dissolved in sterile physiological saline immediately before use. The minimal inhibitory concentrations (MICs) were detected by antimicrobial gradient strip (Etest, bioMérieux, France) according to the manufacturer’s instructions using standard inoculum size; i.e. McFarland standard of 0.5, corresponding to > 10^8^ CFU/ml.

### Batch culture experiments *(in vitro)*

For *in vitro* experiments, bacteria were grown in Lysogeny Broth (LB) as previously described (3). Antibiotics were added to each batch culture at either maximal bacterial growth rate (i.e. at 4 hours of incubation) or at minimal bacterial growth rate (i.e. at 8 hours of incubation), respectively. Samples for quantification of bacterial count, qPCR analysis and fluorescence microscopy were withdrawn pre-treatment (i.e. at 4 or 8 hours of incubation, respectively) and after two hours of antibiotic exposure (i.e. at 6 or 10 hours of incubation, respectively). All samples were immediately set on ice after withdrawal. Control cultures without antibiotic treatment exposure were included in every experiment.

Both treatment studies were performed in duplicate, including both ATCC 25922 wild-type and the genetically modified ALO 4783, in three independent experiments. Both versions of the strain were tested in parallel to ensure the absence of any alterations in growth or antibiotic treatment effect attributable to the transgene insertions. As both growth curves and *ori:ter* were similar for both versions of the strain, these results were pooled for statistical analyses.

Regarding Gentamicin treatment, only the wild-type ATCC 25922 was applied, as the Gentamicin MIC was affected by the presence of a non-removable Kanamycin (KAN) cassette (encoding Kanamycin phosphotransferase) used as clonal selection marker in ALO 4783.

### Mouse peritonitis model experiments *(in vivo)*

The mouse peritonitis model was carried out as previously described, using outbred female NMRI mice (weight 28 ±2 g; Taconic, Denmark) (3). Animals were kept in cages in groups of three; each cage constituting one experimental unit that would be randomly assigned to treatment (CRO, CIP or GEN) or no treatment (control; CTR). Antibiotics were administered as a single bolus injection subcutaneously (s.c.) at either maximal bacterial growth rate (i.e. at 2 hours of infection) or minimal bacterial growth rate (i.e. at 8 hours of infection). Sample collections (peritoneal lavage fluid (PLF), blood, spleen and kidneys) were performed pre-treatment (i.e. at 2 or 8 hours of infection, respectively) in control groups, and after two hours of antibiotic exposure (i.e. at 4 or 10 hours of infection, respectively) in both treatment and control groups. Euthanasia and harvesting of biological specimens were carried out as previously described (3). All biological specimens were immediately placed in an insulated 4°C cooling box for transportation and kept on ice at 4°C until application in subsequent tests. Temporary storage of *E. coli* cultures on ice has previously been demonstrated not to induce any alteration in *in situ* bacterial growth parameters (cell size, *oriC*/cell or *ori:ter*) post harvesting (3).

The mouse peritonitis model was repeated in 4 independent experiments, including a total of 54 animals. Data from repeated experiments were pooled for statistical analyses.

ALO 4783 was applied in all experiments, except from the Gentamicin treatment experiment where ATCC 25922 wild-type was used as infecting agent due to the altered Gentamicin MIC in ALO 4783, as before mentioned.

### Ethics statement

All animal experiments were approved by the Danish Animal Experimentation Inspectorate (licence no. 2014-15-0201-00171) and performed according to institutional guidelines. The mice were regularly observed and scored for signs of distress. Humane end points were constituted by signs of irreversible sickness; the mice would be euthanized upon presentation of any of these signs.

### Quantification of antibacterial activity

Bacterial count measurements from *in vitro* and *in vivo* experiments were performed as previously described (3). Antibacterial activity was measured as the difference between bacterial count pre and post therapy (Δlog_10_ CFU/ml). For meaningful comparison between identical treatment regimens administered at different growth rates (i.e. different pre-treatment bacterial load), antibacterial activity was illustrated as Δlog_10_ CFU/ml relative to the pre-treatment bacterial count (relative Δlog_10_ CFU/ml).

### Quantitative real-time PCR (qPCR)

*ori:ter* was calculated as the population mean level of qPCR amplified *oriC* relative to *terC* from purified bacterial DNA, as previously described (3).

### Fluorescence microscopy

Fluorescence microscopy was used to verify the qPCR data, whenever possible, by direct observation of live single-cell analysis of ALO 4783 carrying fluorescent markers corresponding to the *oriC* and *terC* sites. Fluorescence microscopy analysis was carried out as previously described (3). For Gentamicin treatment experiments, only cell size and morphology were analysed, as the ATCC 25922 wild-type strain was applied. Live bacterial cells were isolated at each sampling time point (i.e. 4, 6, 8 and 10 hours of incubation in the *in vitro* experiments, and 2, 4, 8 and 10 hours of infection in the *in vivo* experiments, respectively). We have previously shown that PLF and blood bacterial population growth in this *in vivo* model do not differ (3). Hence, isolated bacteria from PLF and blood were pooled for analysis. We aimed at isolating 500 bacterial cells at each time point, both with and without antibiotic exposure, but this was not always possible due to substantial or total bacterial clearance in many of the treatment groups, as annotated in Figures 2 and 4.

### Statistical analyses

Bacterial count data were log_10_ transformed prior to analysis. D’Agostino and Pearson omnibus normality test was applied to all data sets. In general, the control group bacterial counts and qPCR data sets represented a normal distribution; those of the treatment groups did not. Statistical significance was evaluated by unpaired *t* test in parametric data and by Mann-Whitney *U* test in nonparametric data. A two-tailed *P-*value of < 0.05 was considered significant. GraphPad Prism version 7 (GraphPad Software, CA, USA) was applied for statistical analysis and illustration.

## Acknowledgements

We thank Jytte Mark Andersen and colleagues at Statens Serum Institut for technical assistance with the animal experiments. The study was partly funded by an EU-IMI Joint Undertaking, grant agreement No. 115583 (ENABLE), by the Scandinavian Society for Antimicrobial Chemotherapy Foundation and by the Danish National Research Foundation (DNRF120) through the Centre for Bacterial Stress Response and Persistence (BASP).

The authors declare no competing interests.

## References

1. Kopf SH, Sessions AL, Cowley ES, Reyes C, Van Sambeek L, Hu Y, Orphan VJ, Kato R, Newman DK. 2016. Trace incorporation of heavy water reveals slow and heterogeneous pathogen growth rates in cystic fibrosis sputum. Proc Natl Acad Sci U S A 113:E110–E116.

2. Olm MR, Brown CT, Brooks B, Firek B, Baker R, Burstein D, Soenjoyo K, Thomas BC, Morowitz M, Banfield JF. 2017. Identical bacterial populations colonize premature infant gut, skin, and oral microbiomes and exhibit different in situ growth rates. Genome Res 27:601–612.

3. Haugan MS, Charbon G, Frimodt-Møller N, Løbner-Olesen A. 2018. Chromosome replication as a measure of bacterial growth rate during Escherichia coli infection in the mouse peritonitis model. Sci Rep 8.

4. Lee AJ, Wang S, Meredith HR, Zhuang B, Dai Z, You L. 2018. Robust, linear correlations between growth rates and β-lactam–mediated lysis rates. Proc Natl Acad Sci 115:4069–4074.

5. Tuomanen E, Cozens R, Tosch W, Zak O, Tomasz A. 1986. The rate of killing of Escherichia coli byβ-lactam antibiotics is strictly proportional to the rate of bacterial growth. Microbiology 132:1297–1304.

6. Brown MR, Collier PJ, Gilbert P. 1990. Influence of growth rate on susceptibility to antimicrobial agents: modification of the cell envelope and batch and continuous culture studies. Antimicrob Agents Chemother 34:1623–1628.

7. Cozens RM, Tuomanen E, Tosch W, Zak O, Suter J, Tomasz A. 1986. Evaluation of the bactericidal activity of beta-lactam antibiotics on slowly growing bacteria cultured in the chemostat. Antimicrob Agents Chemother 29:797–802.

8. Eng RH, Padberg FT, Smith SM, Tan EN, Cherubin CE. 1991. Bactericidal effects of antibiotics on slowly growing and nongrowing bacteria. Antimicrob Agents Chemother 35:1824–1828.

9. Fantin B, Leggett J, Ebert S, Craig WA. 1991. Correlation between in vitro and in vivo activity of antimicrobial agents against gram-negative bacilli in a murine infection model. Antimicrob Agents Chemother 35:1413–1422.

10. Zeiler HJ, Voigt WH. 1987. Efficacy of ciprofloxacin in stationary-phase bacteria in vivo. Am J Med 82:87–90.

11. Zeiler HJ. 1985. Evaluation of the in vitro bactericidal action of ciprofloxacin on cells of Escherichia coli in the logarithmic and stationary phases of growth. Antimicrob Agents Chemother 28:524–527.

12. Korem T, Zeevi D, Suez J, Weinberger A, Avnit-Sagi T, Pompan-Lotan M, Matot E, Jona G, Harmelin A, Cohen N, Sirota-Madi A, Thaiss CA, Pevsner-Fischer M, Sorek R, Xavier RJ, Elinav E, Segal E. 2015. Growth dynamics of gut microbiota in health and disease inferred from single metagenomic samples. Science 349:1101–1106.

13. Brown CT, Olm MR, Thomas BC, Banfield JF. 2016. Measurement of bacterial replication rates in microbial communities. Nat Biotechnol 34:1256–1263.

14. Cooper S, Helmstetter CE. 1968. Chromosome replication and the division cycle of Escherichia coli Br. Am J Med 31:519–540.

15. Wang JD, Levin PA. 2009. Metabolism, cell growth and the bacterial cell cycle. Nat Rev Microbiol 7:822–827.

16. Donachie WD. 1968. Relationship between Cell Size and Time of Initiation of DNA Replication. Nature 219:1077–1079.

17. Helmstetter CE, Cooper S. 1968. DNA synthesis during the division cycle of rapidly growing Escherichia coli Br. J Mol Biol 31:507–518.

18. Hill NS, Kadoya R, Chattoraj DK, Levin PA. 2012. Cell Size and the Initiation of DNA Replication in Bacteria. PLoS Genet 8.

19. Skarstad K, Boye E, Steen HB. 1986. Timing of initiation of chromosome replication in individual Escherichia coli cells. EMBO J 5:1711–1717.

20. Bremer H, Churchward G. 1977. An examination of the Cooper-Helmstetter theory of DNA replication in bacteria and its underlying assumptions. J Theor Biol 69:645–654.

21. Kohanski MA, Dwyer DJ, Collins JJ. 2010. How antibiotics kill bacteria: from targets to networks. Nat Rev Microbiol 8:423.

22. Sezonov G, Joseleau-Petit D, D’Ari R. 2007. Escherichia coli Physiology in Luria-Bertani Broth. J Bacteriol 189:8746–8749.

23. Christensen S, Ladefoged K, Frimodt-Møller N. 1997. Experience with Once Daily Dosing of Gentamicin: Considerations Regarding Dosing and Monitoring. Chemotherapy 43:442–450.

24. Jakobsen L, Cattoir V, Jensen KS, Hammerum AM, Nordmann P, Frimodt-Møller N. 2012. Impact of low-level fluoroquinolone resistance genes qnrA1, qnrB19 and qnrS1 on ciprofloxacin treatment of isogenic Escherichia coli strains in a murine urinary tract infection model. J Antimicrob Chemother 67:2438–2444.

25. Frimodt-Møller N, Bentzon MW, Thomsen VF. 1986. Experimental Infection with Streptococcus pneumoniae in Mice: Correlation of in vitro Activity and Pharmacokinetic Parameters with in vivo Effect for 14 Cephalosporins. J Infect Dis 154:511–517.

26. Knudsen JD, Frimodt-Møller N, Espersen F. 1995. Experimental Streptococcus pneumoniae infection in mice for studying correlation of in vitro and in vivo activities of penicillin against pneumococci with various susceptibilities to penicillin. Antimicrob Agents Chemother 39:1253–1258.

27. Espersen F, Frimodt-Møller N, Corneliussen L, Riber U, Rosdahl VT, Skinhøj P. 1994. Effect of treatment with methicillin and gentamicin in a new experimental mouse model of foreign body infection. Antimicrob Agents Chemother 38:2047–2053.

28. Frimodt-Møller N, Frølund Thomsen V. 1987. The pneumococcus and the mouse-protection test: correlation of in vitro and in vivo activity for beta-lactam antibiotics, vancomycin, erythromycin and gentamicin. Acta Path Microbiol Immunol Scand Sect B 95:159–165.

29. Stouf M, Meile J-C, Cornet F. 2013. FtsK actively segregates sister chromosomes in Escherichia coli. Proc Natl Acad Sci 110:11157–11162.

30. Fontana R. 1998. Interaction of ceftriaxone with penicillin-binding proteins of Escherichia coli in the presence of human serum albumin. J Antimicrob Chemother 42:95–98.

31. Drlica K, Malik M, Kerns RJ, Zhao X. 2008. Quinolone-Mediated Bacterial Death. Antimicrob Agents Chemother 52:385–392.

32. Soriano F, Santamaría M, Ponte C, Castilla C, Fernández-Roblas R. 1988. In vivo significance of the inoculum effect of antibiotics onEscherichia coli. Eur J Clin Microbiol Infect Dis 7:410–412.

33. Nielsen HJ, Hansen FG. 2010. An automated and highly efficient method for counting and measuring fluorescent foci in rod-shaped bacteria. J Microsc 239:194–199.

